# Ligand binding represses bacterial histidine kinase activity by inhibiting its dimerization

**DOI:** 10.1101/2025.05.18.654591

**Authors:** Gaurav D. Sankhe, Jiawei Xing, Merissa Xiao, John Buglino, Huilin Li, Igor B. Zhulin, Michael S. Glickman

## Abstract

Two component systems (TCS) mediate bacterial signal transduction in response to specific environmental conditions. The two components are the sensor kinase (SK), which senses the signal and autophosphorylates on a histidine residue, and a response regulator (RR), which is phosphorylated by the kinase and modifies gene expression. Despite intensive study, the mechanisms of signal sensing by sensor kinases are incompletely defined and the mechanisms by which SKs can sense multiple ligands are unclear. *Mycobacterium tuberculosis* PdtaS/PdtaR is a soluble TCS pair that participates in the Rip1 signal transduction cascade to control virulence by responding to copper and nitric oxide (NO). In contrast to paradigmatic ligand activated SKs, PdtaS is constitutively active without ligand and directly inhibited by Cu or NO, yet it remains unclear how such chemically diverse ligands are sensed. Here we show that PdtaS is a dimeric kinase that constitutively autophosphorylates in trans. Cu and NO both inhibit PdtaS phosphorylation by inhibiting dimerization. Phylogenetic analysis of the PdtaS family reveals conservation of the GAF/PAS dimer interface rather than the ligand binding pockets and mutations in the GAF dimer interface that alter dimerization impair multi-ligand sensing both in vitro and in *M. tuberculosis* cells. These results indicate that a single bacterial kinase can sense chemically diverse inputs through inhibition of dimerization dependent phosphorylation.

## Introduction

Across all domains of life, organisms must sense and respond their external or cellular environments by altering cell state through gene expression changes or altered protein activity. At the most basic level, cellular signaling networks mediate adaptation by sensing diverse conditions and converting this input into an appropriately specific or broad adaptive response [1]. In the simplest case, a signaling ligand binds to a sensory protein and enhances or inhibits its activity, leading to a chain of downstream reactions. However, the mechanisms by which ligand binding changes signaling protein behavior are diverse and still incompletely understood. Ligand binding can affect output response through conformational changes that enhance enzymatic activity, changes in localization, or altered oligomeric state [2, 3]. An additional general design challenge of signaling systems is balancing specificity with receptor diversity. The number of conditions to be sensed is vast, yet encoding monolithic signaling systems that sense only one ligand or condition requires large numbers of germline encoded receptors or a combinatorial system, as occurs in the mammalian immune system. Another solution is to encode signaling systems that can sense multiple ligands which co-occur in a specific condition, but such diversity of sensing must be reconciled with the constraints of protein-ligand interactions and the cost of lost specificity.

All bacteria, both environmental and pathogenic, must adapt to their environments by sensing and responding via modified gene expression [4]. Although there are many mechanisms by which bacteria sense the environment, two component systems (TCS) are a widespread mode in bacterial signal transduction. In the best characterized examples, a sensor kinase (SK), often membrane bound, senses a signal and autophosphorylates on a conserved histidine and then transfers the phosphoryl group to a cytosolic protein known as response regulator (RR). RRs effect the cellular response often through DNA binding and less commonly through second messenger, RNA and protein mediated mechanisms [5–7]. SKs contain a characteristic kinase core consisting of a dimerization histidine phosphotransfer domain (DHp) and an ATP/ADP- binding catalytic domain (CA or C_ATPAse) [8]. Depending upon features of the helix bundle loops between the DHp and CA domains, the autophosphorylation of SKs can occur in cis (intramolecular) or trans (intermolecular)[9]. When SKs phosphorylate in *trans* [10–13] the ATP bound to the CA domain of one SK provides the phosphoryl group to the histidine residue of the dimeric partner. Autophosphorylation in trans therefore necessitates dimerization for kinase activity [8, 14]. Cis autophosphorylation occurs when the ATP bound to the CA domain phosphorylates the histidine residue on the same SK polypeptide [15–17].

SKs sense a diverse spectrum of physical and chemical cues such as light [18], temperature [19], pH [20], NO [21], quinones from the respiratory chain [22] cyclic dinucleotides [23], cations [24] and many other compounds. SKs are usually activated by ligand-induced conformational changes leading to kinase activation and autophosphorylation. However, for many TCSs, the ligands that directly bind and activate the SK remain elusive [25]. In contrast to ligand activation, in some cases ligand binding inhibits signal transduction, as observed for WalK upon Zn^2+^ binding to the PAS domain [24] and the inhibition of the autophosphorylation of FixL by oxygen [26].

*M. tuberculosis* encodes 13 two-component systems, most of which follow the classic cognate sensor kinase-response regulator pairing, often encoded in an operonic arrangement in the chromosome [27, 28]. These include the intensely studied DosS/R system [21, 29], the PhoP/R system [20, 30, 31] and the essential MtrAB system [32–34]. The PdtaS/PdtaR TCS is atypical in several ways. First, PdtaS (encoded by *rv3220*) and PdtaR (encoded by *rv1626*) are encoded at distinct sites in the chromosome. The PdtaS kinase has a C terminal kinase domain and N terminal GAF and PAS domains, classically involved in ligand binding [35–38]. However, studies of PdtaS kinase activity indicate that PdtaS is constitutively active without ligands [39–41]. In addition, the PdtaR RR is not a DNA binding transcription factor. In addition to its receiver domain, which is phosphorylated by PdtaS, PdtaR contains an ANTAR domain, which binds RNA hairpins [42]. Although the full functions of ANTAR domain RNA binding have not been defined, in some cases the ANTAR domain interaction regulates transcriptional termination [43, 44].

PdtaS and PdtaR were implicated in the Rip1 pathway of *M. tuberculosis* signal transduction by a genetic suppressor screen in which inactivation of either PdtaS or PdtaR reverted the copper and nitric oxide sensitivity of *M. tuberculosis* lacking *rip1*. Copper and NO directly inhibit the kinase activity of PdtaS, an inhibition that requires the N terminal GAF and PAS domains [39], indicating that the GAF-PAS are necessary to transmit the inhibitory signal to the kinase domain. The suppressor mutations in PdtaS were both in the N terminal GAF domain. One was a premature termination mutation, but the other was a single amino acid substitution (V54F) in the GAF domain near a dicysteine motif (C53/C57). Mutation of C53 or C57 to alanine did not affect constitutive PdtaS kinase activity but did abolish PdtaS inhibition by both copper and NO [39].

These data indicate that PdtaS is a direct sensor of multiple ligands, including NO and Cu, and that inhibition of autophosphorylation is a mechanism of PdtaS signaling. However, it remains unclear how a single kinase can sense such chemically diverse ligands. Ligand binding pockets of GAF and PAS domains can bind a wide variety of ligands[38], but it remains to be determined whether multi-ligand sensing by PdtaS represents a manifestation of specific chemical recognition by the GAF-PAS domains or some other mechanism.

In this work, we demonstrate that PdtaS is a constitutively active SK which autophosphorylates in *trans*, thereby necessitating kinase dimerization for activity. Cu and NO both inhibit PdtaS phosphorylation by inhibiting dimerization. Phylogenetic analysis of the PdtaS family reveals conservation of the GAF/PAS dimer interface but not the ligand binding pockets. Mutations in the GAF dimer interface that alter dimerization also impair multi-ligand sensing of Cu and NO *in vitro* and in *M. tuberculosis* cells. Our findings establish a mechanism of multi ligand sensing through alteration of sensor oligomeric state.

## Results

### PdtaS phosphorylates as a dimer in trans

Although many TCS are dimeric, the mechanism of autophosphorylation by sensor kinases can be either in cis or trans. Cis autophosphorylation occurs when ATP bound by a single kinase polypeptide transfers phosphate to the histidine on the same molecule, whereas trans autophosphorylation occurs when ATP bound by one kinase monomer transfers phosphate to the histidine on the dimeric partner. To investigate the autophosphorylation mechanism of PdtaS, we generated a structural model of the full length PdtaS dimer, including C terminal kinase, GAF and PAS domains (Figure 1A). Detailed examination of the ATP binding site revealed close proximity to the phosphorylated histidine 303 on the opposite PdtaS kinase domain in the dimer, possibly suggesting trans autophosphorylation (Figure 1B). To test which mode of phosphorylation is operative in PdtaS, we mixed wild type PdtaS with PdtaS-H303Q, which can bind ATP but cannot be phosphorylated due to a mutation in the acceptor histidine, and measured autophosphorylation using [γ−^32^P] ATP. Increasing PdtaS-H303Q inhibited the phosphorylation of wild type PdtaS, a result that suggests trans phosphorylation since cis phosphorylation should not be inhibited by an inactive dimeric partner (Fig 1C). A similar inhibition of PdtaS autophosphorylation was observed with titration of the inactive and ATP binding defective PdtaS-G443A, which in turn gained weak phosphorylation from the dimeric WT protein (Figure 1D). To confirm trans phosphorylation, we mixed PdtaS-H303Q with PdtaS-G443A and quantitated autophosphorylation. Each protein is inactive either due to loss of ATP binding or the phosphoacceptor site (Fig 1C,D), but a mixture of the two proteins restored phosphorylation of G443A, consistent with trans autophosphorylation of G443A with ATP binding proficient H303Q­­ (Fig 1E,F). To confirm that the inactivity of each mutant protein is not due to loss of dimerization, we measured the dimerization affinity of PdtaS by microscale thermophoresis (MST) with itself, PdtaS-H303Q or PdtaS-G443A and found nearly identical dimerization affinities (Fig 1G, WT-WT-1400nM, WT-H303Q-1443nM, WT-G443A-1457nM). These results demonstrate that PdtaS constitutively autophosphorylates in trans.

**Figure 1:**
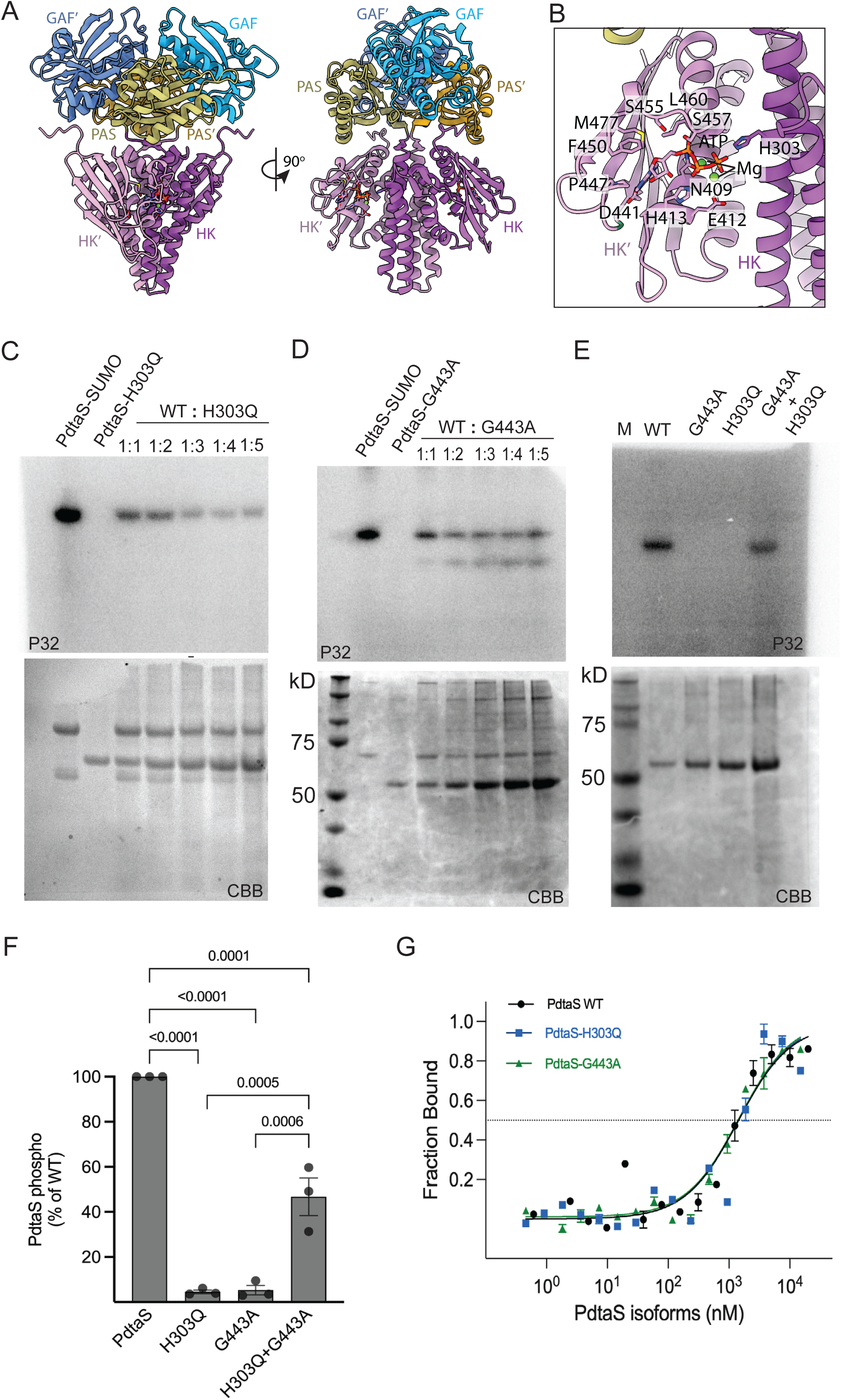
PdtaS phosphorylates in trans. **A.** Cartoon representation of the AlphaFold3 predicted structures of the PdtaS dimer with ATP and Mg2^+^ bound in two orthogonal views. The two PdtaS monomers are distinguished by intensity of color shading and individual domains with (‘). **B.** Expanded view of the dimeric HK domain, showing the potential trans-phosphorylation model. Residues interacting with ATP are shown as sticks. G443 is highlighted in green. **C.** Autophosphorylation assay showing autoradiogram (P32, upper panel) or protein (Coomassie brilliant blue, CBB, lower panel) of SDS PAGE gels from kinase reactions containing gamma ^32^P. The reactions contain PdtaS-SUMO alone, PdtaS-H303Q, which is inactive, or mixtures of these two proteins with escalating concentrations of H303Q. **D.** Autophosphorylation assay with the same design as panel (C) using ATP binding defective PdtaS-G443A **E.** Trans-autophosphorylation assay demonstrates phosphorylation in trans. Autophosphorylation as described in panel (D) of WT PdtaS, PdtaS-G443A, PdtaS-H303Q, or a 1:1 mixture of G443A and H303Q. Each mutant protein is inactive, but PdtaS-H303Q can phosphorylate G443A in trans. **F.** Quantitation of biologic replicates of the assay in (E) with WT PdtaS normalized to 100%. P values were calculated by one way ANOVA with Tukey’s multiple comparison correction and error bars are SEM. **G.** Dimerization affinities measured by MST using PdtaS, PdtaS-H303Q, or PdtaS-G443A each measured against WT PdtaS. Error bars are SEM, and each point represents average of 3 replicates

### Ligands that inhibit PdtaS activity inhibit dimerization

Having established an autophosphorylation assay that specifically reflects PdtaS dimerization, we tested the ability of Cu and NO to inhibit autophosphorylation. Although 10 μM calcium had no effect, 10 μM Cu strongly inhibited PdtaS trans autophosphorylation (Fig 2A). Similarly, 100 μM spermine NONOate strongly inhibited trans autophosphorylation in comparison to spent NONOate control (Fig 2A). Although these data indicate that PdtaS ligands inhibit trans phosphorylation, as demonstrated previously with wild type PdtaS protein [39], this effect could either occur by inhibition of phosphotransfer without dissociation of the PdtaS dimer, or by ligand induced dissociation of the PdtaS dimer. To test whether Cu and NO modify dimer formation, we quantitated the affinity of dimer formation by MST using titration of unlabeled PdtaS with fluorophore conjugated PdtaS. We detected PdtaS dimer formation with an apparent Kd of 1543 nM (Fig 2B, 2D). Addition of 10μM Nickel or calcium did not change the apparent Kd, but addition of 10μM Cu inhibited dimerization with an apparent Kd in the presence of Cu of 5072 nM (p=0.0088 for buffer vs Cu by one way ANOVA, Fig 2B, 2D). NO also inhibited dimerization, increasing the apparent Kd from 1196 nM for spent NO control to 5717nM with 100μM NO (p=0.0044 by one way ANOVA, Fig 2C, 2D). These data indicate that both Cu and NO specifically inhibit PdtaS dimer formation at concentrations that inhibit trans autophosphorylation.

**Figure 2:**
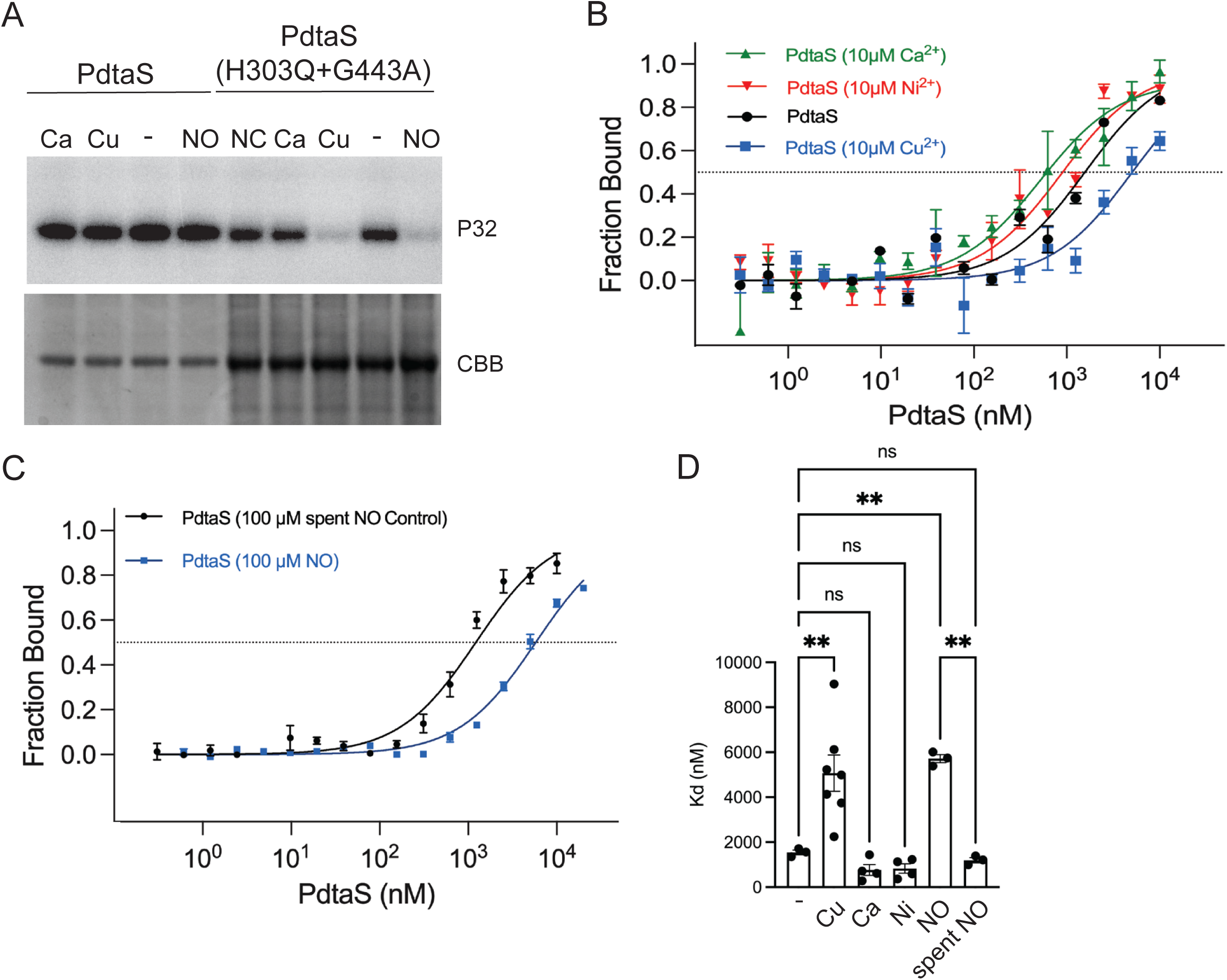
Ligands that inhibit PdtaS activity inhibit dimerization. **A.** Autophosphorylation assay with either wild type PdtaS or the trans autophosphorylation assay from 1E, with 10 μM Ca, 10 µM Cu, no addition (-), 100 µM spermine NONOate (NO), or 100 µM spent spermine NONOate (NC). **B.** Copper inhibits PdtaS dimerization. Microscale thermophoresis (MST) was used to measure the dimerization affinity of PdtaS prelabeled with amine coupling dye (RED-NHS 2^nd^ generation Nanotemper Technologies) and unlabeled PdtaS in the presence of buffer (black), 10µM Cu (blue square), 10µM Ni (red triangle), or 10µM Ca (green triangle). Each point is the average of 4 replicates and error bars are SEM. **C.** Nitric oxide inhibits PdtaS dimerization. MST as described in panel B with 100μM spermine NONOate or spent NO control. Each point is average of 3 replicates and error bars are SEM. **D.** Quantitation of dimerization dissociation constant (Kd, nM) calculated from replicate MST experiments in the presence of Cu, Ca, Ni, NO, or spent NO. Statistical significance is indicated and calculated by one way ANOVA with Bonferroni correction for multiple comparisons.

### Bioinformatic analysis of the PdtaS kinase family reveals conservation of the dimer interface

Several biochemical properties of the PdtaS kinase differ from the usual model of sensor kinase activation. Most prominent is the constitutive kinase activity, which is present without any activating ligand. Although cyclic di-GMP has been proposed as a ligand for the PdtaS GAF domain, this ligand is not needed for kinase activity [41]. Additionally, copper and NO, two other ligands for PdtaS, inhibit kinase activity and inhibit dimerization, possibly suggesting that dimerization could be a mechanism of ligand sensing. The GAF and PAS domains that are N terminal to the PdtaS kinase domain (Figure 1A) are widespread protein domains in signal transduction systems with diverse functions, including direct ligand binding and dimerization [38, 45]. Specifically, based on our previous study, the PAS domains of PdtaS homologs form a cluster with a distinct conservation pattern [38](Supplementary Fig. 1). To expand this analysis, we collected 4,988 homologs from the reference protein sequence database using *M. tuberculosis* PdtaS as the query, which resulted in 887 sequences after removing redundancy (Supplementary Dataset 1, see Methods). We performed bioinformatic analyses of the conservation of PdtaS across the Actinomycetota phyla and mapped this conservation onto the predicted full length PdtaS dimer structure predicted using AlphaFold3 [46], which agrees well with the truncated experimentally determined structure (PDB accession 2YKF, see Figure S4 for model quality) [35]. We found that conserved residues (>50% conservation) were distributed along the dimer interface in GAF, PAS, and kinase domains (Fig 3A, Supplementary Dataset 2). In comparing the number of conserved residues along the GAF and PAS dimer interfaces to the surface of the GAF/PAS ligand binding cavity, we found more conservation (>90%) along the dimer interface than in the cavities, particularly for the PAS domain (Fig 3B, Supplementary Dataset 2). This analysis suggests that the PdtaS kinase family has evolved to conserve the dimerization interface, shown above to be important for autokinase activity, but that the putative ligand binding domains do not have a conserved ligand cavity, arguing against a single specific ligand that binds in the GAF or PAS pocket in this family of histidine kinases. Although cyclic-di-GMP (cDG) was reported to stimulate kinase activity of PdtaS and affect its behavior in vivo, we did not observe an effect of cDG on the dose response of inhibition of PdtaS by copper (Figure S3).

**Figure 3.**
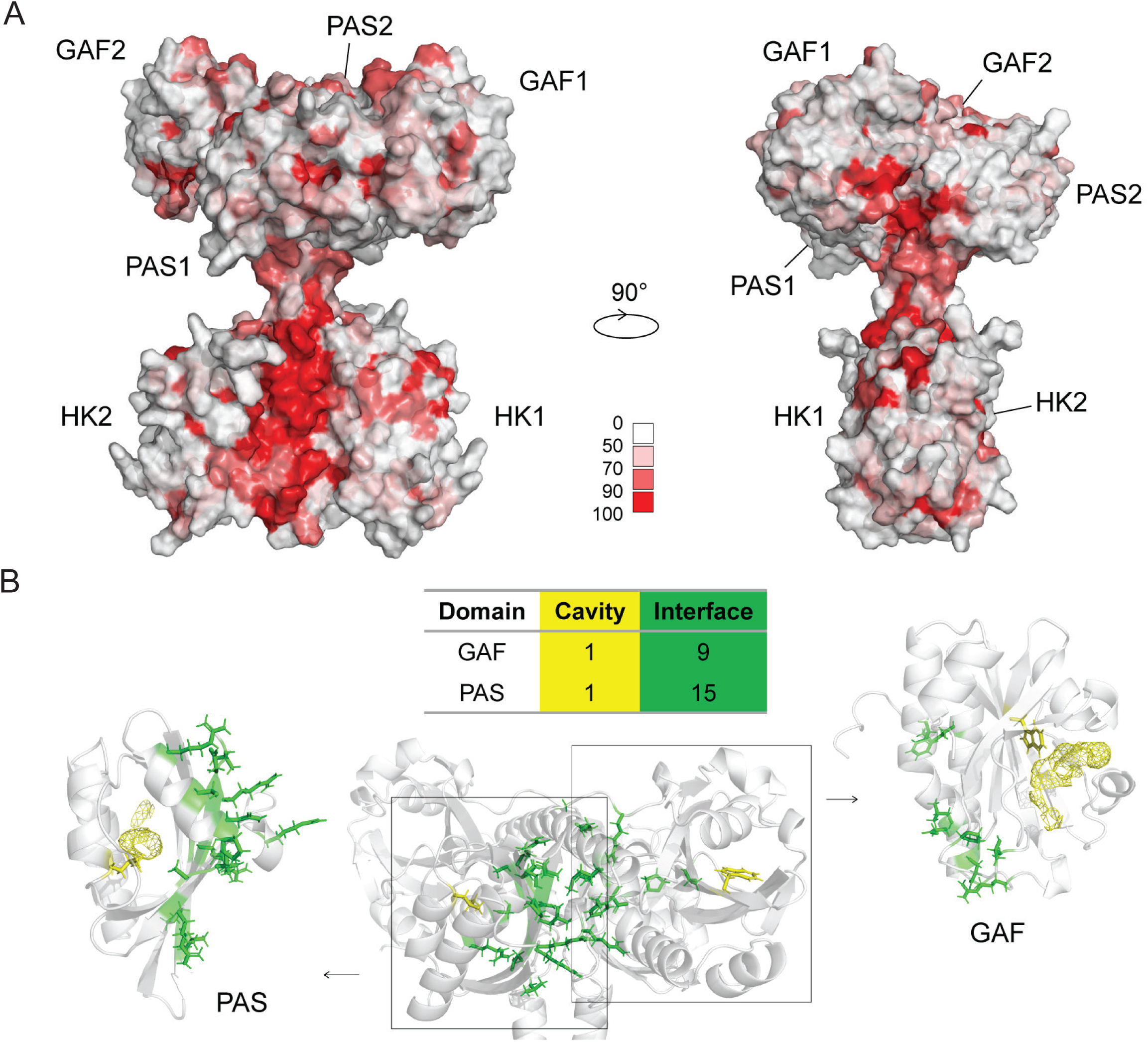
Conservation of the Dimer interface of PdtaS. **A.** Structural model of the PdtaS dimer with PAS, GAF, and Histidine Kinase (HK) from each monomer labeled. The conservation across Actinomycetota PdtaS proteins is indicated by red shading at different conservation levels. **B.** Interface residues of the GAF and PAS domains are more conserved than ligand­­­­ pocket residues. The table displays the number of conserved amino acid residues (present in >90% sequences) across Actinomycetota PdtaS orthologs in the ligand binding cavity (yellow) or dimerization interface (green). The structural model shows the GAF and PAS domain of one PdtaS monomer in the dimer structure and separated in an exploded view with color coding of residues according to the scheme in the table with interface residues in green and cavity residues in yellow.

### The cysteine motifs in the PdtaS GAF domain modulate ligand sensing and dimer formation

Our prior data indicated that a dicysteine motif (C53/C57) in the N terminal PdtaS GAF domain (Fig 4A) was critical to ligand sensing. PdtaS-C53A or -C57A were both active kinases but could not be inhibited by Cu or NO [39]. However, the mechanism by which these cysteines mediate ligand mediated kinase inhibition was not clear. We hypothesized that, given the effect of Cu and NO on PdtaS dimerization, the dicysteine motif might alter the dimer kinetics as a mechanism of ligand insensitivity. We first queried the conservation of C53 and C57 across PdtaS orthologs. We found that although C57 is conserved among a minority of PdtaS proteins in the Actinomycetales, the Mycobacteriales order had striking conservation of these residues. Almost all PdtaS proteins in the Mycobacteriales had conservation of C53 or both C53 and C57, with a minority with only C57 (Fig 4B, Supplementary Dataset 1). To test whether C53 or C57 affect PdtaS dimerization, we purified PdtaS-C53A and PdtaS-C57A proteins and measured dimerization Kd by MST. In comparison to WT PdtaS, both cysteine mutants displayed enhanced dimerization affinities (WT:1318nm, C53A:445nM, C57A: 608nM, WT vs C53A p<0.001, WT vs C57A p=0.008, both by one way ANOVA with Bonferroni correction, Fig 4C,D). These data suggested that the conserved cysteines in the GAF weaken the dimer, and that resistance of the GAF domain mutants to ligand inhibition might be due to enhanced dimer formation. To test this hypothesis, we measured dimerization Kd of the C53A (Figure 4E) and C57A (Figure 4F) proteins by MST in the presence of Cu and NO. In contrast to the wild-type protein, the dimerization affinities of both cysteine GAF mutants were unaffected by either Cu or NO. These data indicate that the ligand sensing by the GAF domain of PdtaS is mediated through dimerization affinity.

**Figure 4:**
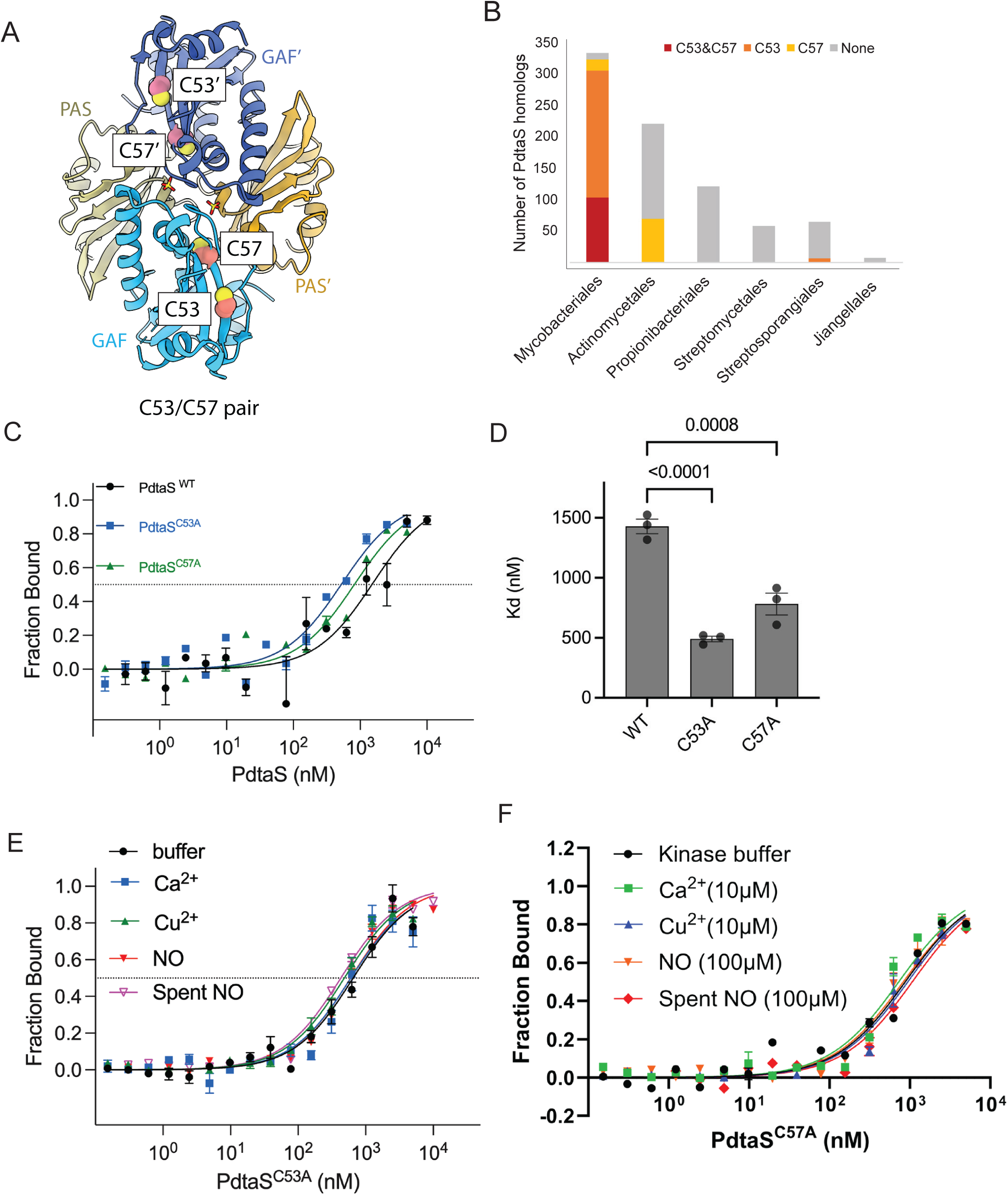
A dicysteine motif in the GAF domain modifies dimerization affinity and multiligand sensing. **A.** Close-up view of the C53/C57 dicysteine motif in the GAF-PAS dimer structure (PDB ID: 2YKF). C53 and C57 are presented as spheres. **B.** Conservation of C53 and C57 in PdtaS according to bacterial order **C.** Effects of mutations at C53 or C57 on dimerization. Shown are MST plots of wild type (black), PdtaS C53A-C53A (blue), or PdtaSC57A-C57A (green) **D.** Quantitation of dimerization affinities of the indicated PdtaS proteins by MST of biologic triplicates. P values were calculated by one way ANOVA with Bonferroni correction and error bars are SEM. **E,F** NO and copper do not inhibit PdtaSC53A-C53A (E) or PdtaSC57A-C57A (F) dimerization. MST curves measuring the dimerization affinity of the indicated PdtaS proteins in the presence of the indicated ligands.

### Dimerization separation of function mutations impair PdtaS kinase activity

In analyzing the dimer interface of PdtaS, we identified H67 and R137 as conserved residues positioned in the dimer interface of the GAF domains (Fig 5A), suggesting that they may be involved in dimerization mediated ligand sensing. We changed each residue to alanine by site directed mutagenesis and purified PdtaS-H67A and PdtaS-R137A. PdtaS-R137A was refractory to purification due to insolubility, but PdtaS-H67A was purified successfully. We tested the dimerization affinity of PdtaS-H67A in comparison to PdtaS and observed reduced dimerization with a Kd of 8777nM for H67A and 1501 nM for wild type (Fig 5B). Autophosphorylation assays indicated that PdtaS-H67A was ∼3 fold less active than wild type kinase (Fig 5C), consistent with impaired dimerization controlling PdtaS kinase activity.

**Figure 5:**
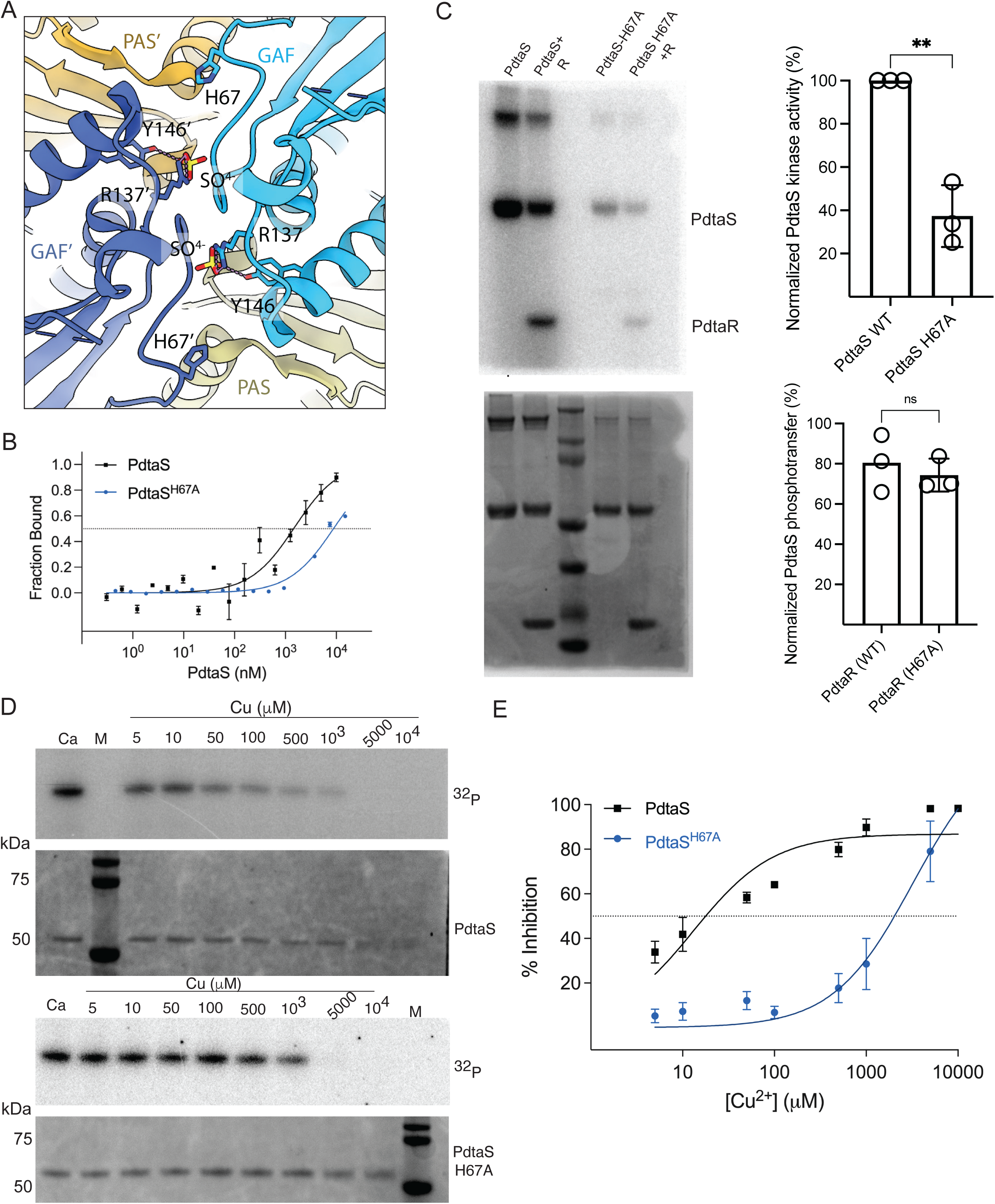
GAF-PAS dimer interface controls ligand inhibition of PdtaS. **A.** The GAF-PAS dimer interface (PDB ID: 2YKF). H67 and R137 are in sticks. R137 and Y146 coordinate a sulfate in the crystal structure. Hydrogen bonds are labeled as dashed lines. **B.** PdtaSH67A-H67A has a weakened dimer interface. MST measurement of the dimerization affinities of WT PdtaS or PdtaSH67A-H67A. **C.** The GAF dimer interface affects autophosphorylation but not phosphotransfer. Autophosphorylation (PdtaS alone) or phosphotransfer reactions (PdtaS+PdtaR) with autoradiograms (^32^P, top) or Coomassie stained gels (CBB, bottom) for either WT PdtaS or PdtaS-H67A. The graphs to the right quantitate autophosphorylation or normalized phosphotransfer to PdtaR. **D.** Copper does not inhibit PdtaS-H67A. Autophosphorylation reactions as in Figure 1 of WT or PdtaS-H67A with increasing concentrations of copper as indicated and Calcium (Ca) as a negative control. The dose response curve is derived from quantitation of biologic triplicates. **E.** Copper inhibition curves from triplicate experiments as in (D) WT PdtaS (black) or PdtaS H67A. Error bars are SEM and the curves are significantly different by ANOVA (p<0.01) at Cu concentrations 5-1000μM.

However, incubation with PdtaR revealed that both wild type and H67A kinases were capable of phosphotransfer to PdtaR. Although the absolute quantity of PdtaR phosphorylated by PdtaS-H67A was reduced, normalization of the PdtaR intensity to the PdtaS autophosphorylated intensity revealed that phosphotransfer was unaffected by the H67A mutation (Fig 5C), indicating that this mutation specifically affects autophosphorylation but not phosphotransfer.

To test whether the impairment of PdtaS dimerization affects ligand inhibition by Cu, we tested PdtaS autophosphorylation in the presence of escalating concentrations of Cu. Although wild type PdtaS was inhibited, as previously observed[39], PdtaS-H67A was highly resistant to Cu inhibition with a Ki of 3262 mM vs 13.14 for wild type (Fig 5D,E). These data indicate that weakening the dimer affinity of PdtaS dampens the dimer dependent trans autophosphorylation activity and prevents further ligand inhibition.

### Inhibition of PdtaS through GAF-PAS interdomain coupling

The data above indicates that the PdtaS kinase dimerizes in part through contacts in the N terminal GAF domain. GAF mutations that either strengthen or weaken dimerization have corresponding effects of kinase activity and inhibition by ligands that inhibit PdtaS autokinase activity. However, the kinase domain at which ATP binding and autophosphorylation occur is C terminal, suggesting that there must be some mechanism to couple the inhibition of dimerization to the kinase domain. We further analyzed the PdtaS conservation and interpreted conserved residues considering the predicted interdomain structure of the kinase dimer. Position 28 of the GAF domain is occupied by aspartate or glutamate in 100% of PdtaS proteins in the Actinomycetota phyla (Fig 6A, B, Supplementary Dataset 2). E28 of *M. tuberculosis* PdtaS is predicted to form an interdomain contact to arginine 261 of the PAS domain (Fig 6A,B), which is also conserved in 99% of PdtaS proteins. We hypothesized that this interdomain connection might be involved in coupling the GAF domain dimer dissociation signal to the C terminal PAS to ultimately inhibit autophosphorylation. The dimerization affinity of PdtaS-R261A by MST was identical to WT, indicating that the GAF-PAS bridge does not affect dimerization (Fig 6C). In contrast to wild type PdtaS, for which Cu inhibits dimerization, copper had no effect on the dimer affinity of the PtdaS-R261A protein in comparison to calcium or control conditions (Kd WT=1656 vs R261A=1480, Fig 6D). However, the GAF-PAS bridge mutant (R261A) displayed an attenuated inhibition by Cu in comparison to wild type (Fig 6E, WT Ki 10.52 µM vs PdtaS-R261A Ki 181.9 µM). These results are consistent with a model in which GAF-PAS interdomain coupling transmits the dimer dissociation signal to inhibit phosphorylation.

**Figure 6.**
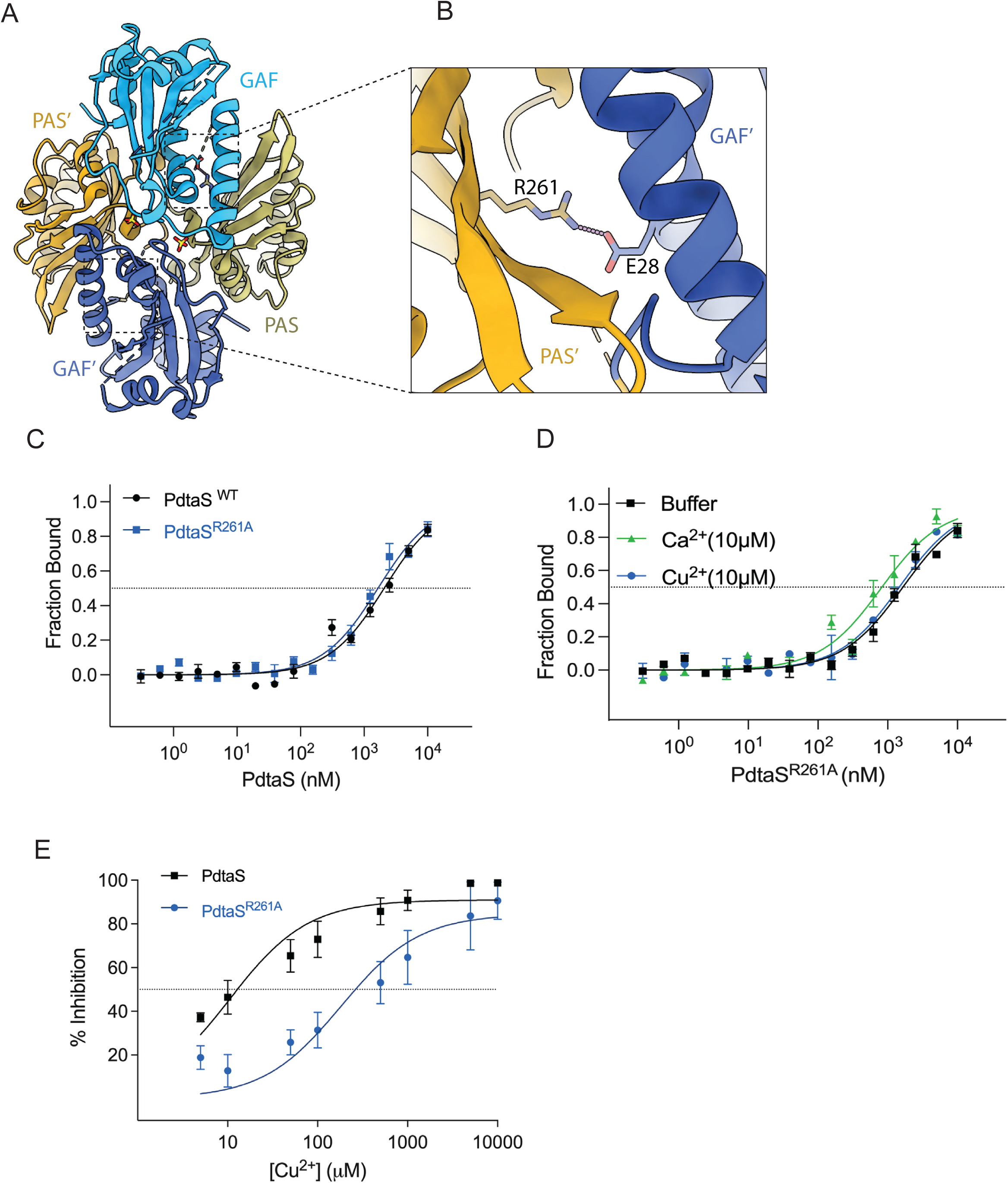
Interdomain coupling of the GAF dimerization signal. **A.** Cartoon view of the GAF-PAS dimer (PDB ID 2YKF). **B.** Close-up views of the interaction of E28 and R261 at the interface of GAF and PAS domain. **C.** Interdomain bridge does not affect dimerization. MST dimerization curves of the indicated proteins. **D.** The interdomain bridge is required for ligand induced inhibition of dimerization. MST curves of PdtaS-R261A with buffer, calcium, or copper, demonstrating no effect on apparent dimer affinity. **E.** The interdomain bridge is required for ligand inhibited autophosphorylation. Inhibition curves of autophosphorylation assays of WT PdtaS or PdtaS-R261A with increasing copper concentrations.

### PdtaS dimerization mediates signaling in vivo

To test whether PdtaS dimerization is important for PdtaS signaling in vivo, we expressed PdtaS alleles carrying mutations that affect dimerization (C53, C57, or H67) or interdomain coupling (R261), as defined above. Inactivating mutations in PdtaS suppress the copper sensitivity of the *M. tuberculosis* Δ*rip1* strain [39], providing a sensitive system to interrogate PdtaS mutants. Complementation of *M. tuberculosis* Δ*rip1pdtaS-*F37X with PdtaS alleles encoding an ALFA epitope tag allowed monitoring of PdtaS expression (Fig 7A). All PdtaS protein were detectable, although all mutant proteins accumulated to lower levels than WT PdtaS (Fig 7A). We tested the copper sensitivity of these strains and observed that complementation with PdtaS-ALFA restored the Cu sensitivity to Δ*rip1pdtaS*F37X, indicating that ALFA tag does not impair PdtaS function. In contrast, PdtaS-C53, PdtaS-C53A/C57A and H67A were inactive. However, PdtaS-R261A, despite its resistance to Cu inhibition in vitro (Figure 6), was still active in vivo (Figure 7B). Although the three dimerization mutants that are inactive are expressed at lower levels than WT PdtaS, their expression levels are similar to R261A (Figure 7A), indicating that these expression levels are sufficient for signaling.

**Figure 7.**
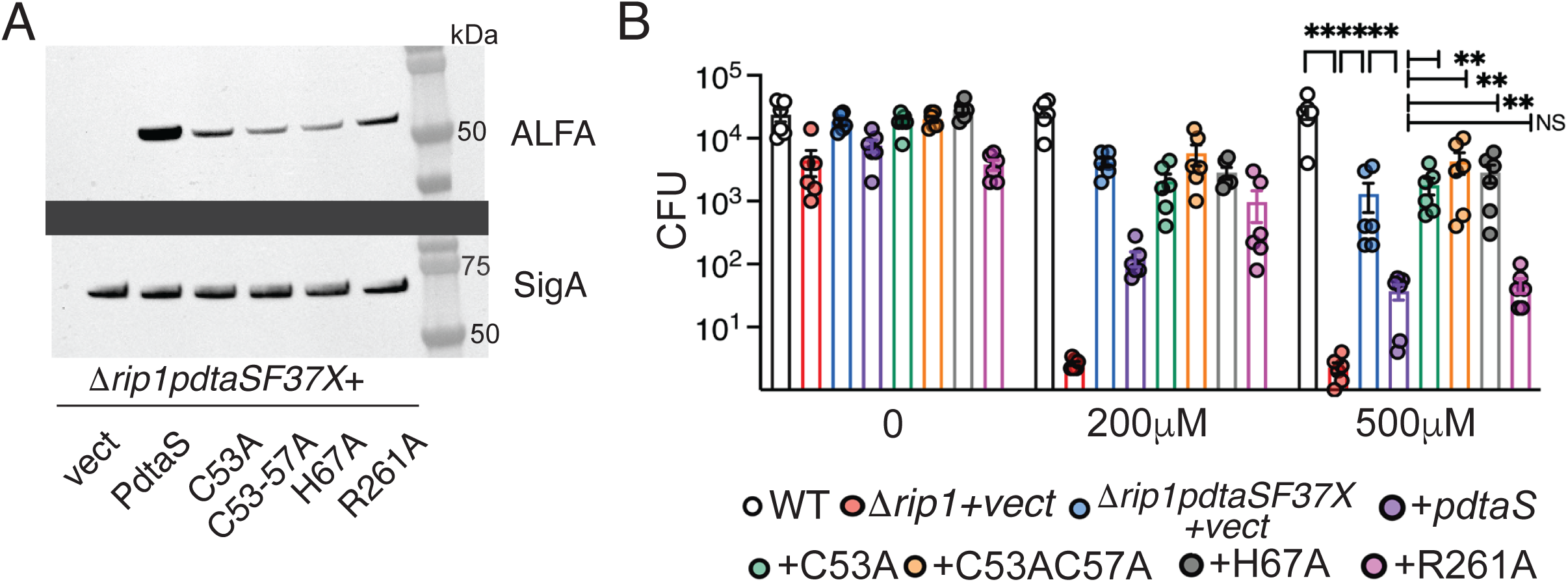
In vivo function of the PdtaS dimerization interface in copper resistance. **A.** Immunoblot of lysates from *M. tuberculosis* strains with anti-ALFA nanobody (top) or anti-SigA antibodies (bottom) as a loading control. Strains are all derived from *M. tuberculosis* Erdman Δ*rip1pdtaS* F37X, which carries a frameshift suppressor mutation in PdtaS as described in [39]. Each strain carries an ALFA tagged PdtaS complementation plasmid encoding nothing (vect), wild type PdtaS, or PdtaS with the indicated mutations **B.** Copper sensitivity of *M. tuberculosis* expressing PdtaS dimerization mutant proteins. The *M. tuberculosis* strains in panel A, along with wild type and Δ*rip1* strain were cultured on agar media containing no addition or 200/500 μM copper sulfate. Each point is a biologic replicate from two technical replicates and statistical significance for the 500 μM condition by the Mann-Whitney test with correction for multiple comparisons. Error bars are SEM. **=p<0.01.

## Discussion

We have investigated the mechanism of kinase activation of the *M. tuberculosis* PdtaS kinase. PdtaS was previously implicated in sensing copper and nitric oxide as part of the Rip1 signal transduction cascade, in addition to binding cyclic di-GMP [39, 41]. PdtaS is a soluble cytoplasmic protein that constitutively autophosphorylates in vitro without any added ligand. This constitutive activity is inhibited by both copper and nitric oxide and this inhibition shuts off the PdtaS/PdtaR pathway, thereby modifying gene expression via the RNA binding activity of the PdtaR ANTAR domain [39]. Although dimeric sensor kinases can autophosphorylate either in cis or trans [9], our data indicate that PdtaS phosphorylates in trans. The paradigmatic mechanism of signal transduction by sensor kinase-response regulator pairs is that a signal is sensed by the SK through specific ligand recognition, activating the kinase to autophosphorylate and then phosphorylate the response regulator. PdtaS works by an inverted regulatory logic in which chemically diverse ligands inhibit the constitutively active kinase, but molecular and mechanistic insights for its sensing mechanism were unknown.

We propose that inhibition of dimerization is the mechanism by which PdtaS can sense chemically diverse ligands rather than by specific recognition by the GAF and PAS domains, which are usually specific ligand sensors. This model is based on the following data:

1. Copper and NO inhibit dimer formation by PdtaS
2. Sequence analysis of the *M. tuberculosis* PdtaS protein indicates that mycobacterial PdtaS belongs to a distinct cluster. Sequence conservation of this cluster is observed in the dimer interface of the GAF and PAS domains rather than in the putative ligand binding pockets.
3. Two cysteine residues in the GAF domain, C53 and C57, are conserved in Mycobacteriales PdtaS. When mutated to alanine, PdtaS forms a tighter dimer, and these cysteine mutant proteins are resistant to inhibition by both Cu and NO.
4. Histidine 67 is at the dimer interface and PdtaS-H67A forms a weaker dimer than WT PdtaS but is fully competent for phosphotransfer to PdtaR. PdtaS H67A is resistant to inhibition by Cu.
5. PdtaS E28-R261 are predicted to form a bridge between the GAF and PAS domains, suggesting that they might transmit a signal between these two domains. Although PdtaS-R261A dimerizes with equal affinity as WT PdtaS, Cu cannot dissociate the dimer and therefore inhibition of autokinase activity by Cu is attenuated.
6. All of the mutations at the dimerization interface that abolish Cu sensing in vitro similarly inactivate PdtaS function in in vivo in controlling Cu resistance. However, the R261 mutation, which abolishes Cu sensing in vitro, is nevertheless active in vivo, suggesting that this interdomain coupling model is incomplete or compensated by other mechanism in vivo to inhibit trans autophosphorylation. However, this R261A PdtaS protein serves as a useful control for the expression levels of the dimerization mutants. Although all mutant proteins are underexpressed compared to WT PdtaS, R261A is still active and expressed at similar levels to the other proteins, suggesting the lack of activity of dimerization mutant proteins is not due to expression level.

Taken together, these data are consistent with a model in which modulation of dimer affinity is the sensing mechanism of the mycobacterial clade of PdtaS kinases, rather than specific recognition of Cu or NO by the ligand binding pockets of the GAF or PAS domains. We propose that this mechanism of dimer sensing allows integration of multiple inputs into the kinase without the constraints of specific ligand recognition. It is also possible that this mechanism is operative in a cytoplasmic kinase for which dimer association is in solution rather than being constrained by the membrane, as is the case with membrane bound SKs. There are other examples of SKs recognizing multiple ligands in Salmonella [47, 48], possibly suggesting that this mechanism could extend to other pathogens.

Although we identify mutations with both positive and negative effects on dimer affinity which have effects of ligand inhibition of the kinase, this data does not identify specific molecular details on how dimerization is inhibited and whether Cu and NO both interact with the same regions of the dimer interface. Although it is possible that NO could covalently modify the cysteines we identify as conserved and important for signaling, our data does not determine if PdtaS cysteines are directly modified. It is also possible that other factors present in the bacterial cell might modify the dimerization affinity and thereby tune the setpoint of the PdtaS/PdtaR system similar to setting the signal detection thresholds of various mycobacterial SKs through sequestration by non-cognate RRs [49]. A recent global phosphoproteomic study, for example, [50] identified multiple phosphorylation sites in PdtaS which could modify the responses observed in vitro. Overall, our data reveal a new mechanism of SK sensing based on dimer affinity, a mechanism that may be applicable to other SKs for which the sensing mechanisms are not well defined.

## Materials and Methods

### Reagents

Antibiotics IPTG (isopropyl β-D-1-thiogalactopyranoside) and DTT (dithiothreitol) were purchased from GoldBio Inc., USA. Glycine, ATP, Tween-20 from Merck Sigma Aldrich, USA. Protein marker from Thermo Scientific, USA. Co^2+^-NTA TALON metal affinity resin from Takara, Japan. Protease inhibitor cocktail from Amresco and γ^32^P labelled ATP (>3000 Ci/mmol) from Perkin Elmer, USA. Media and salts were purchased from Fisher Scientific, USA.

### Recombinant PdtaS/PdtaS mutant protein purification

Full length C-terminal 10x His tagged PdtaS/PdtaS mutant, and N-terminal 10xHisSmt3 PdtaS/PdtaS mutant fusion constructs, were both expressed and purified (Figure S2) from Rosetta 2 (DE3) *E.coli* (Millipore Sigma). Cultures, 2-4L, grown in Luria-Bertani (LB) media to an OD_600_ of 0.8 at 37°C were equilibrated to 30°C for 30 min prior to induction. Protein expression was then induced with 0.5 mM isopropyl-β-D-thiogalactoside (IPTG), and the cultures were allowed to grow post induction for 3-4 additional hours at 30°C. Cells were then collected by centrifugation, re-suspended in lysis (350mM sodium chloride, 20mM Tris-HCl pH 8.0, 20mM imidazole, 0.5mM β-mercaptoethanol) at a ratio of 2mL of buffer to 1g of wet pellet weight and stored at -80°C. The following day frozen cell pellets were thawed and disrupted by 2 passes thought a French pressure cell at an internal pressure of 14000 psi (Sim-Aminco, 40000 psi cell). Crude lysates were clarified by centrifugation at 25,000xg for 40 minutes in an SS-34 Rotor (Sorvall), and then passed over TALON metal affinity resin (Takara, Japan) equilibrated in wash buffer. Columns were then washed with 20 column volumes of Wash buffer (350mM sodium chloride, 20mM Tris-HCl pH 8.0, 40mM imidazole, 0.5mM β-mercaptoethanol). Proteins bound to the resin were eluted with 2 column volumes of Elution Buffer (Wash Buffer + 200mM imidazole).

After elution from the TALON column PdtaS 10XHis was dialyzed and concentrated to ∼2mg/ml in a dialysis buffer containing 50mM Tris-HCl pH 8.0, 50 mM sodium chloride, 100 µM DTT (dithiothreitol) with 40 %(v/v) glycerol.

Eluted 10xHis-Smt3-PdtaS/PdtaS mutant protein was further purified as follows. 10XHis-Smt3 was cleaved from PdtaS/PdtaS mutant protein by overnight treatment with Ulp1 protease at 4°C. Free PdtaS was then recovered in the flow through fraction of a second round of TALON resin purification. PdtaS/PdtaS mutant protein was then dialyzed and stored in the dialysis buffer.

### PdtaS mutagenesis

Site-directed mutagenesis was performed in PdtaS expression vectors were using the QuikChange^©^ site-directed mutagenesis protocol using Q5^®^ Hi-Fidelity DNA polymerase (New England Biolabs, USA) and primers synthesized having the desired point mutations. The presence of mutations in the plasmids was confirmed using DNA sequencing and proteins were purified as described above.

### Sensor Kinase autophosphorylation activity using PAGE/Autoradiography technique

5 μM of the purified sensor kinase (SK) PdtaS/PdtaS mutant protein was incubated for 20 minutes in the autophosphorylation buffer (50 mM Tris HCl at pH 8.0, 50 mM KCl, 10 mM MgCl_2_) containing 50 μM ATP and 1μCi of γ^32^P-labelled ATP at 30°C. The reaction was terminated by adding 1× SDS-PAGE sample buffer (2% w/v SDS, 50 mM Tris HCl, pH 6.8, 0.02% w/v bromophenol blue, 1% v/v β-mercaptoethanol, 10% v/v glycerol). The samples were resolved on 15% v/v SDS-PAGE. After electrophoresis the gel was washed and exposed to phosphor screen (Fujifilm Bas cassette2, Japan) for 4 hours followed by imaging with Typhoon 9210 phosphorimager (GE Healthcare, USA). Quantitative densitometric analysis of the bands present in autoradiogram corresponding to respective proteins visualized in Coomassie brilliant blue (CBB) staining, was carried using ImageJ software.

### Sensor kinase mechanism of activation (cis or trans)

PdtaS-H303Q [41] or PdtaS-G443A [51] were co incubated in the indicated escalating concentrations with a fixed amount of 5 μM of PdtaS wild type protein for 10 mins in the autophosphorylation buffer at 30°C. The reaction was initiated by adding 50 μM and 1 μCi of γ^32^P-labelled ATP for 20 mins at 30°C and terminated by adding 1X SDS-PAGE sample buffer and separation by PAGE/Autoradiography. Similarly, the effect on kinase activity of 5 μM of PdtaS protein was observed by titrating with increasing amounts of ATP binding defective mutant (mutation in the catalytic ATPase domain),. 5 μM each of purified PdtaS^H303Q^ and PdtaS^G443A^ mutant proteins were co-incubated in the autophosphorylation buffer for 10 mins at 30°C. The reaction was started by adding 50 μM of unlabeled and 1 μCi of γ^32^P-labelled ATP at 30°C for 20 mins and terminated by adding 1X SDS-PAGE sample buffer. Samples were then processed using PAGE/Autoradiography technique as described above.

### PdtaS autophosphorylation inhibition assays

5 μM of the purified SK PdtaS was pre incubated for 10 minutes in the autophosphorylation buffer with copper chloride at various concentrations as indicated in the figure legend and with 1 mM of calcium chloride as a control. Similarly, the effect nitric oxide (NO) and was studied by preincubating 5 μM of PdtaS protein with spermine NONOate (Cayman Chemical Company, USA) as a rapid NO donor for 40 minutes at a 100 μM concentration. A negative control for spermine NONOate was prepared by preincubating 5 μM of PdtaS with 100 μM of spermine NONOate which had been allowed to exhaust NO release by overnight incubation at room temperature in Tris HCl buffer at pH= 6.8. The autophosphorylation reaction was initiated by adding 50 μM ATP along with 1μCi of γ^32^P-labelled ATP at 30°C for 20 minutes. The reaction was terminated by adding 1× SDS-PAGE sample buffer. The samples were resolved on 15% v/v SDS-PAGE. After electrophoresis the gel was washed and exposed to phosphor screen (Fujifilm Bas cassette2, Japan) for 4 hours followed by imaging with Typhoon 9210 phosphorimager (GE Healthcare, USA). Quantitative densitometric analysis of the bands present in autoradiogram corresponding to respective proteins visualized in Coomassie brilliant blue (CBB) staining was carried using ImageJ software. The dose response curves were plotted, and inhibition constant (*K_i_*) was obtained by fitting the percentage inhibition to concentration plot using one site specific binding fit in Graphpad Prism.

### Determination of binding affinity of PdtaS G443A or H303Q by MicroScaleThermophoresis (MST)

Purified PdtaS protein (100 nM) was prelabeled using amine coupling Monolith Protein Labeling kit RED-NHS 2^nd^ generation (Catalog No. MO-L011, Nanotemper Technologies, GmbH) as prescribed in manufacturer’s protocol. This fluorescently labeled PdtaS was mixed with increasing concentrations of wild type PdtaS (0.61 nM to 10 μM) or mutant PdtaS^H303Q^ (0.45 nM to 15 μM) or mutant PdtaS^G443A^ (0.45 nM to 15 μM) in the autophosphorylation buffer with 50 μM of ATP and 0.025% Tween-20 for 10 mins kept at 30°C. The sample was then loaded into standard treated capillaries using a Monolith NT. 115 instrument (NanoTemper Technologies GmbH). The red laser was used for a duration of 35 s for excitation (MST power = 40%, LED power 40%). The data were analyzed using MO Control software (NanoTemper Technologies GmbH) to determine the apparent *K_D_* values represented as fraction bound.

### Determination of binding affinity for dimerization using MST

100 nM of PdtaS was prelabeled using amine coupling Monolith Protein Labeling kit RED-NHS 2^nd^ generation (Catalog No. MO-L011, Nanotemper Technologies, GmbH) as prescribed in manufacturer’s protocol. To determine binding affinity for dimerization this fluorescently labeled PdtaS protein was mixed with increased concentration of PdtaS protein (0.61 nM to 10 μM) in the autophosphorylation buffer having 0.025% Tween-20 and 50 μM of ATP for 10 mins at 30°C. Similarly, binding affinity for dimerization of PdtaS^C53A^, PdtaS^C57A^, PdtaS^R261A^ and PdtaS^H67A^ was determined using MST by pre labeling them with the amine coupling RED-NHS 2^nd^ generation dye as prescribed and mixing the same respective unlabeled proteins in increasing concentration of 0.15 nM to 5 μM for PdtaS C53A and C57A, 0.3 nM to 2.5 μM for PdtaS R261A and 0.45 nM to 15 μM for PdtaS H67A respectively. Effect of individual ligands and their respective controls on dimerization of PdtaS or its mutant were similarly measured using MST by preincubating the ligand at concentrations mentioned in the respective figure legends.

### Sequence collection and conservation analysis

4,988 homologous sequences of PdtaS were collected from the NCBI reference protein sequence database using a default BLAST search with *M. tuberculosis* PdtaS WP_003416886.1 as the query [52]. Sequence redundancy was reduced at a 90% level using CD-HIT, resulting in 887 sequences[53]. These sequences were aligned using MAFFT with the E-INS-i algorithm[54]. Sequence conservation was calculated using a custom python script based on the percentage of each residue in the multiple sequence alignment. Conserved residues at the interfaces were identified by PyMOL by selecting residues within 4.0 Å of other domains[55]. Conserved residues at the cavities were identified by CASTp with a default parameter[56]. Protein structures were visualized using PyMOL[55]. Bacterial taxonomy was determined using the Genome Taxonomy database (GTDB)[57].

### Strain construction, verification, and Cu sensitivity testing

*M.tuberculosis pdtaS* ALFA point mutant strains were generated by overlap extension PCR and verified by sanger sequencing of the resulting plasmids. *M. tuberculosis* Erdman Δ*rip1pdtaS* F37X competent cells were transformed with the verified plasmids and the resulting transformants were confirmed by immunoblotting and tested for resistance to copper toxicity as previously described [39].

## Acknowledgements

This work was supported by grant R01AI138446 (to MSG), P30CA008748, R35GM131760 (to I.B.Z), R01AI070285 (to HL), and a Van Andel Institute P2i grant (to MX).

## Supplementary Data

**Figure S1**

Sequence conservation of the PdtaS PAS domain cluster 120 from Xing et al. [38]

**Figure S2**

Purified proteins used in this study, Coomassie stained SDS-PAGE gel of the specified PdtaS proteins used in biochemical assays.

**Figure S3**

Effect of c-di-GMP on PdtaS kinase activity.

**(A)** Autophosphorylation assays of PdtaS in the presence of 10μM calcium, escalating concentrations of Cu, 0 or 1000 μM cyclic-di-GMP, and escalating concentrations of Cu with a fixed concentration of 100 μM cyclic-di-GMP. The top panel shows the autoradiogram and the bottom gel shows the stained PAGE gel with protein quantities in each lane.

**(B)** Quantitation of autokinase inhibition derived from triplicate experiments as pictured in (A). Error bars are SEM and calculated Ki values derived from the best fit curves as solid lines are indicated.

**Figure S4.**

AlphaFold confidence metrics for the PdtaS dimer. The modeled sequence comprises two identical protomers, with positions 1–501 corresponding to chain A and positions 502–1002 corresponding to chain B.

**(A)** Predicted Aligned Error (PAE) plots for the top five ranked models. The low error regions (blue diagonals) indicate that the individual domains within each protomer are structurally stable. The higher error regions (red off-diagonals) demonstrate high flexibility in the interplay between the PAS/GAF domain and the kinase domain, as well as dynamic interactions between the opposing kinase domains. This reflects the structural rearrangements required for ligand regulation and trans-phosphorylation.

**(B)** 3D structural model of the PdtaS dimer colored by predicted Local Distance Difference Test (pLDDT) scores. High-confidence domains are colored blue (>90), while kinase regions colored red (<50) represent the dynamic structural flexibility required for kinase functioning.

**(C)** Per-position pLDDT scores for all five ranked models, quantitatively illustrating the distribution of stable structural domains and flexible regions across both protomers in the dimer.

**Supplementary Dataset 1**

PtdaS proteins using *M. tuberculosis* PdtaS as a query

**Supplementary Dataset 2**

Conservation of PdtaS amino acids positions

