## Supplementary figures and images for "Ligand binding represses bacterial histidine kinase activity by inhibiting its dimerization"

### Figures S1-S4

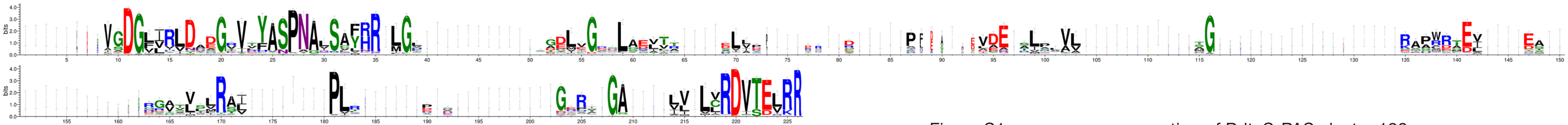

Figure S1: sequence conservation of PdtaS-PAS cluster 120

WebLogo 3.7.4

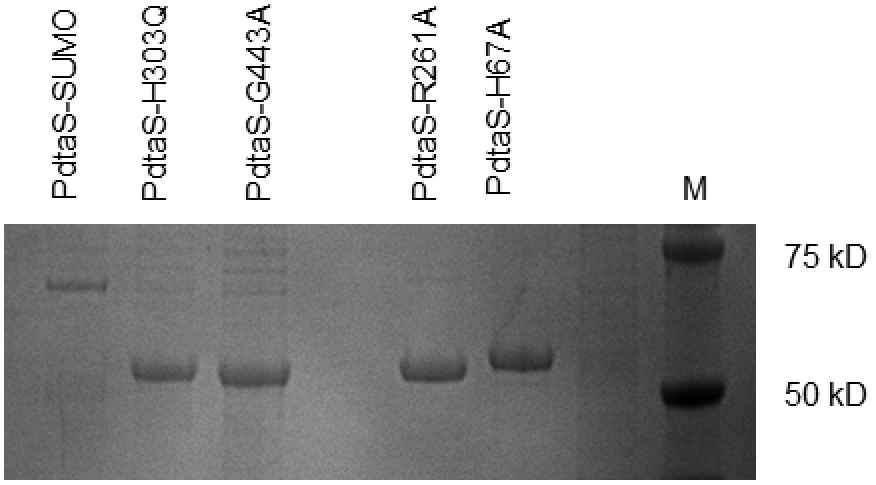


Figure S2: purified proteins

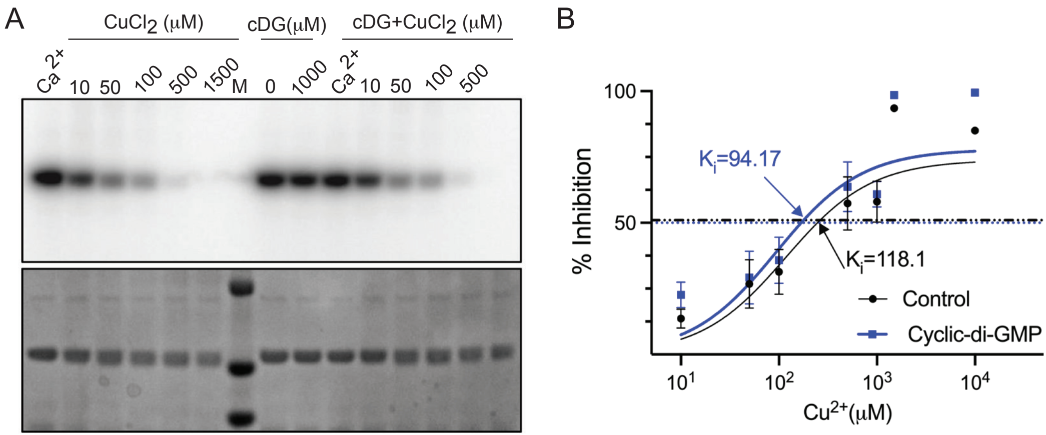

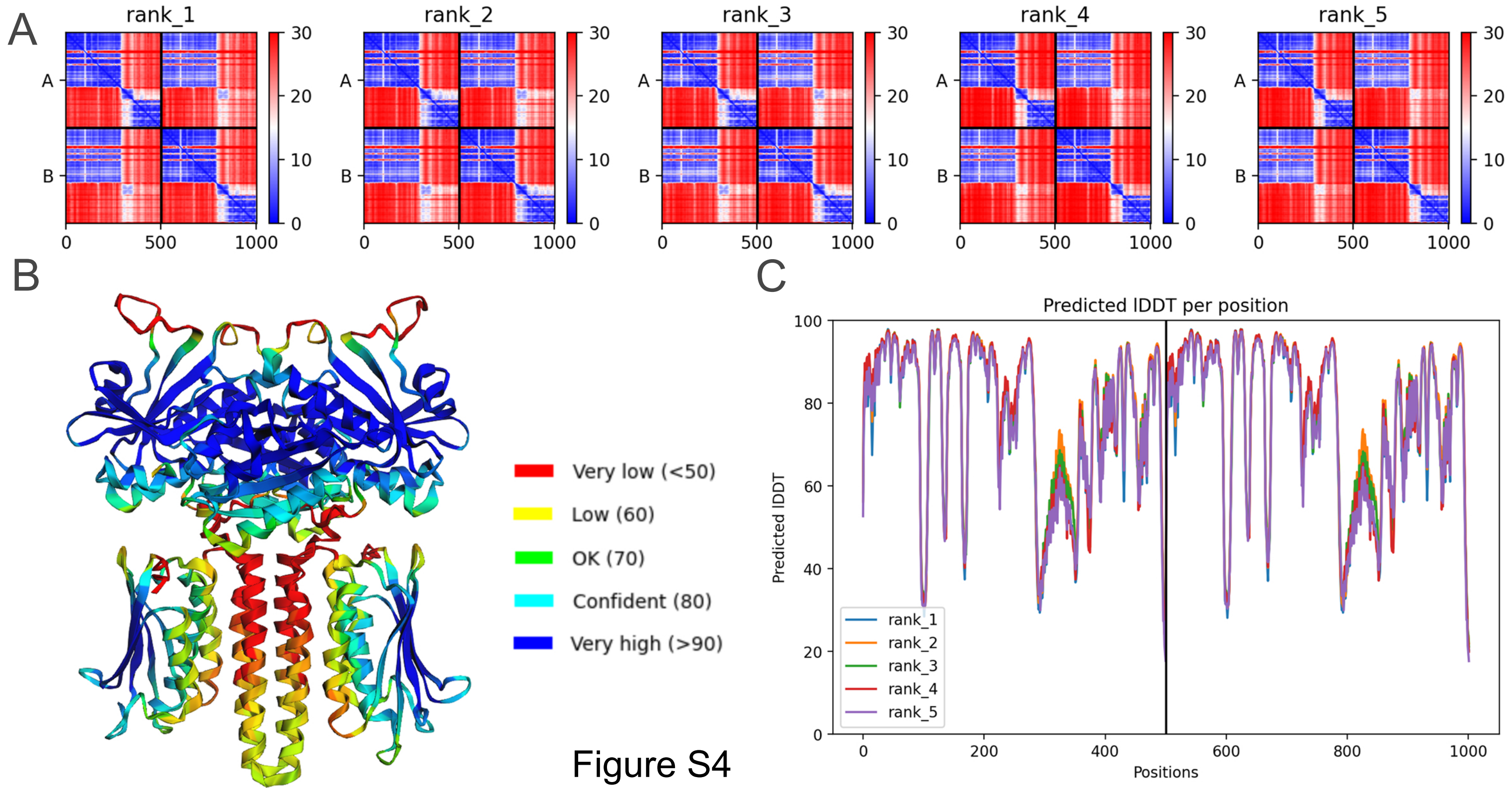
